# Convergent mutations and single nucleotide variants in mitochondrial genomes of modern humans and Neanderthals

**DOI:** 10.1101/190363

**Authors:** Renata C. Ferreira, Camila R. Rodrigues, James R. Broach, Marcelo R. S. Briones

## Abstract

Genetic contributions of Neanderthals to the modern human genome have been evidenced by comparison of present-day human genomes with paleogenomes suggesting that the Neanderthal introgression is higher in Asians and Europeans and lower in Africans. Neanderthal signatures in extant human genomes are attributed to intercrosses between Neanderthals and archaic Anatomically Modern Humans (AMH). Although Neanderthal signatures are well documented in the nuclear genome, it has been proposed that there is no contribution of Neanderthal mitochondrial DNA to contemporary human genomes. Here we show that modern human mitochondrial genomes contain potential 66 Neanderthal signatures, or Neanderthal single nucleotide variants (N-SNVs) being 36 in coding regions of which 7 are nonsynonymous. Also, 7 N-SNVs are associated with traits such as cycling vomiting syndrome, Alzheimer’s disease, Parkinson’s disease and 2 N-SNVs are associated with intelligence quotient. Based on recombination tests, Principal Component Analysis (PCA) and the complete absence of these N-SNVs in 41 archaic AMH mitogenomes we conclude that convergent evolution due to homoplasy and not recombination, explains the presence of N-SNVs in present-day human mitogenomes.

## 1. Introduction

Comparative analyses of present day human genomes and paleogenomes led to the proposal that Neanderthals contributed to the modern makeup of the human genome [1–4]. The current interpretation of genomic signatures is that Neanderthal contributions are different in European, East Asian and African lines of descent, with a higher frequency of Neanderthal segments in Asians and Europeans and lower frequencies in Africans [1]. Intercrosses between Neanderthals and ancient *Homo sapiens* lineages, or archaic Anatomically Modern Humans (AMH) who migrated from Africa into the Middle East and Europe in the last 50,000 years might explain the presence of Neanderthal signatures in extant human genomes [3,4]. The spatio-temporal overlap of Neanderthals and archaic AMH is estimated to be approximately 22,000 years since the first AMH arrived in Europe around 50,000 years ago and the last Neanderthal remains (in Spain) date back to 28,000 years [5,6].

Although there is evidence for Neanderthal contributions to the modern nuclear genome, it has been proposed that there is no Neanderthal contribution to contemporary human mitochondrial genomes (mitogenomes) [7]. It is widely accepted that human mitochondrial DNA is exclusively inherited from the mother and lacks recombination [8]. Because of mitogenome matrilineal inheritance this implies that the intercrosses occurred exclusively between Neanderthal males and AMH females or that crosses between AMH males and Neanderthal females were extremely rare. Another possibility is that crosses between AMH males and Neanderthal females produced such unfavorable trait combinations, due to mitonuclear incompatibility [9], that none of their descendants left marks in present day human populations.

The lack of recombination in human mtDNA is still a matter hotly debated, with published evidence supporting it and rejecting it [10,11]. The main problem posed by a complete lack of human mtDNA recombination is the consequential Müller ratchet, the theoretical concept, in population genetics, that explains the accumulation of harmful mutations in a population over time, which helps to understand genetic deterioration and extinction [12,13]. Although the Müller ratchet is a theoretical concept it highlights the importance of recombination in the long-term maintenance of genetic diversity and the prevention of genetic deterioration in populations. Recombination allows for the removal of harmful mutations and the shuffling of genetic material and therefor alleviate the effect of the Müller ratchet [14].

Analysis of mitogenomes of present-day humans, archaic AMH and Neanderthals revealed 66 variants (N-SNVs), or signatures, in contemporary human mitogenomes that were also present in Neanderthals but not in archaic AMH mitogenomes. The archaic AMH samples selected for analysis are approximately the same age as Neanderthal mitogenomes, and therefore represent, in theory, an archaic AMH ensemble that could have overlapped with Neanderthals in Europe, Middle East and Central Asia. This also allows for assessment of homoplasy because Neanderthal and archaic AMH sequences here considered are of approximately the same age. The distribution of these N-SNVs in different human haplogroups was investigated and Principal Component Analysis (PCA) and recombination tests were performed to evaluate whether recombination or homoplasy might explain the N-SNVs in present human mitogenomes.

## 2. Material and Methods

### 2.1. Mitochondrial genome sequences

Comparative analysis of the mitochondrial DNA from present day humans (52 sequences), ancient *Homo sapiens*, or archaic AMH (41 sequences) and Neanderthals (9 sequences) were aligned to identify single nucleotide variants. Paleogenomes mtDNA sequences were downloaded from GenBank for 41 archaic AMH (*H. sapiens*) and 9 Neanderthal mtDNA (Table 1). Neanderthal signatures, or N-SNVs, were determined as positions that were identical between present day sequences and Neanderthal sequences at the exclusion of archaic AMH. The 52 sequences of present day human mtDNA, representing all major mitochondrial haplogroups (Table 2), were selected from the PhyloTREE database [15] and downloaded from GenBank.

**Table 1.** Archaic AMH and Neanderthal mitogenomes used in this study. Haplogroup inference quality** as calculated by Haplogrep 2. [20].

| *Archaic AMH* | Haplogroup | Quality** (%) | Age (years) | Location | Reference | GenBank/ENA* |
| --- | --- | --- | --- | --- | --- | --- |
| Berry Au Bac 1 | U5b1a | 98.78 | 7,160-7,319 | France | [39] | KU534977 |
| Bockstein | U5b1d1 | 98.72 | 8,016-8,329 | Germany | [39] | KU534973 |
| Cuiry Les Chaudardes 1 | U5b1b | 97.01 | 8,050-8,360 | France | [39] | KU534975 |
| Ofnet | U5b1d1 | 98.72 | 8,159-8,424 | Germany | [39] | KU534974 |
| Felsdach | U5a2c | 97.43 | 8,380-8,980 | Germany | [39] | KU534954 |
| Hohlenstein Stadel | U5b2c1 | 94.77 | 8,446-8,809 | Germany | [39] | KU534979 |
| Falkenstein | U5b2a | 92.19 | 8,993-9,409 | Germany | [39] | KU534980 |
| Mareuil Les Meaux 1 | U5a2+16362 | 100 | 9,080-9,500 | France | [39] | KU534959 |
| Les Closeaux 3 | U5a2 | 98.27 | 9,580-10,230 | France | [39] | KU534958 |
| Ranchot 88 | U5b1 | 95.95 | 9,933-10,235 | France | [39] | KU534978 |
| Ibousseries 31-2 | U5b1+16189 | 99.01 | 10,140 | France | [39] | KU534976 |
| Ibousseries 25-1 | U5b2a | 93.92 | 10,140 | France | [39] | KU534981 |
| Ibousseries 39 | U5b2b | 92.40 | 11,600-12,040 | France | [39] | KU534972 |
| Hohle Fels 10 | U8a | 96.92 | 12,700 | Germany | [39] | KU534961 |
| Rochedane | U5b2b | 97.01 | 12,830-13,090 | France | [39] | KU534971 |
| Burkhardtshohle | U8a | 96.92 | 14,150-15,080 | Germany | [39] | KU534960 |
| Hohle Fels 79 | U8a | 96.92 | 14,270-15,070 | Germany | [39] | KU534962 |
| Brillenhohle | U8a | 97.66 | 14,400-15,120 | Germany | [39] | KU534947 |
| Oberkassel 998 | U5b1+ | 99.45 | 14,000 | Germany | [53] | KC521457 |
| Goyet Q-2 | U8a | 96.92 | 14,780-15,230 | Belgium | [39] | KU534963 |
| Rigney 1 | U2'3'4'7'8'9 | 87.68 | 15,240-15,690 | France | [39] | KU534957 |
| Hohle Fels 49 | U8a | 96.92 | 15,568-16,250 | Germany | [39] | KU534964 |
| Paglicci 71 | U5b2b | 97.97 | 18,197-18,973 | Italy | [39] | KU534950 |
| Dolni Vestonice 43 | U5 | 97.44 | 25,000 | Czech Rep. | [39] | KU534970 |
| Goyet Q56-16 | U2 | 91.40 | 26,040-26,600 | Belgium | [39] | KU534965 |
| Goyet 2878-21 | U5 | 92.05 | 26,269-27,055 | Belgium | [39] | KU534955 |
| Goyet Q376-19 | U2 | 89.43 | 27,310-27,720 | Belgium | [39] | KU534967 |
| Goyet Q55-2 | U2 | 85.86 | 27,310-27,730 | Belgium | [39] | KU534948 |
| La Rochette | M | 92.38 | 27,400-27,784 | France | [39] | KU534951 |
| Goyet Q53-1 | U2 | 93.23 | 27,720-28,230 | Belgium | [39] | KU534966 |
| Paglicci 108 | U2'3'4'7'8'9 | 92.40 | 27,831-28,961 | Italy | [39] | KU534968 |
| Paglicci 133 | U8c | 99.41 | 28,000-29,000 | Italy | [39] | KU534956 |
| Dolni Vestonice 16 | U5 | 97.94 | 29,386-30,567 | Czech Rep. | [39] | KU534949 |
| Doni Vestonice 14 | U5 | 100 | 31,000 | Czech Rep. | [54] | KC521458 |
| Cioclovina 1 | U | 96.72 | 32,519-33,905 | Romania | [39] | KU534969 |
| Goyet Q376-3 | M | 88.67 | 33,140-33,940 | Belgium | [39] | KU534953 |
| Goyet Q116-1 | M | 93.64 | 34,430-35,160 | Belgium | [39] | KU534952 |
| Kostenki 14 | U2 | 92.69 | 36-39,000 | Russia | [53] | FN600416* |
| Fumane 2 | R | 94.43 | 39-41,000 | Italy | [53] | KP718913 |
| Tianyuan | B4'5 | 91.17 | 40,000 | China | [37] | KC417443 |
| Ust-Ishim | R | 87.89 | 45,000 | West Siberia | [53] | PRJEB6622* |
| **Neanderthals** |  |  |  |  |  |  |
| Vindija 33.16 | H1e | 52.74 | 38,000 | Croatia | [42] | AM948965 |
| Vindija 33.17 | H1e | 52.75 | Not dated | Croatia | [55] | KJ533544 |
| Vindija 33.19 | H1e | 52.74 | Not dated | Croatia | [55] | KJ533545 |
| Vindija 33.25 | H1as | 52.84 | Not dated | Croatia | [56] | FM865410 |
| El Sidron 1253 | H1e | 52.81 | 39,000 | Spain | [53,56] | FM865409 |
| Feldhofer 1 | H1as | 52.84 | 40,000 | Germany | [56] | FM865407 |
| Feldhofer 2 | H1e | 52.81 | 40,000 | Germany | [56] | FM865408 |
| Altai Neanderthal | L1'2'3'4'5'6 | 55.96 | 50,000 | Siberia | [3] | KC879692 |
| Mezmaskaya 1 | L1'2'3'4'5'6 | 52.12 | 65,000 | Russia | [56] | FM865411 |

**Table 2.** Haplogroups of present-day *Homo sapiens sapiens* mitogenomes used in this study.

| **Mitochondrial Haplogroup** | **GenBank** | **Mitochondrial Haplogroup** | **GenBank** |
| --- | --- | --- | --- |
| L0a1b1 | AF381988 | S1 | AF346963 |
| L1c3a | AF381992 | Q1 | AY289090 |
| L2a1f | AY195776 | Z1 | AY519493 |
| L3d3b | AF381998 | O | AY289059 |
| L4a1 | FJ460531 | Y1 | KF540727 |
| L5a1a | DQ341060 | R0a | JX153281 |
| L6a | EU092773 | R0a1a3 | GU592021 |
| N1b1a3 | AY195756 | R1a | KC985147 |
| N9a1 | HM589048 | I1 | JQ245776 |
| N2a | JF904935 | W1 | EU558696 |
| A | AP013225 | X1a | EU600318 |
| B2 | EF648602 | X3 | EF177437 |
| F1a1 | NA17963 | U1a1d | EF692533 |
| M29a | DQ137407 | U2c | AY714010 |
| M2b | EU443512 | U6a7a2 | AF382008 |
| M3b | FJ383523 | K | AF382005 |
| M8a1 | KF148510 | K1 | EU073969 |
| M9a | HM346891 | K2a2a | EU327986 |
| M20 | JX289112 | H1a1 | EU007858 |
| G1a1 | HM460792 | H3 | EU150187 |
| E1 | KF540505 | H15 | KC911292 |
| D4 | JQ704974 | J1c | EU547187 |
| C1a | EU007858 | J2a2a | EF660967 |
| C4 | FJ951604 | T1 | JQ797976 |
| C7 | FJ951594 | V1a | JQ702026 |
| P2 | AY289088 | V2 | JQ703647 |

### 2.2. Sequence assembly and alignments

The Ust-Ishim sequence was assembled using reads downloaded from Study PRJEB6622 at the European Nucleotide Archive (EMBL-EBI) and assembled using the CLC Genomics Workbench 7 program (https://www.qiagenbioinformatics.com). To maintain the reference numbering, sequences were aligned to the revised Cambridge Reference Sequence (rRCS; GenBank accession number NC012920) [16], totalizing 103 sequences using the map to reference option implemented in Geneious 10 program [17]. Variants were called using Geneious 10 program. A total of 918 polymorphic positions were found. Neanderthals, ancient and modern humans were screened for disease associations at MitoMap (http://www.mitomap.org/MITOMAP) [18].

### 2.3. Phylogenetic inference

Position specific similarities between modern haplogroups and Neanderthals were depicted by cladograms for each of the single 66 variant positions present only in Neanderthals and modern humans and excluding archaic AMH, were generated using parsimony heuristic search implemented in PAUP v4.1a152 with the default parameters [19]. Proximity of mitochondrial haplogroups in Ancient *H. sapiens* and Neanderthals were inferred using Haplogrep 2.1.0 [20]. All 66 cladograms, corresponding to each N-SNV, are available upon request.

### 2.4. Recombination analysis

Potential recombination between Neanderthals and ancient *H. sapiens* sequences was inferred by a phylogenetic based method implemented by manual bootscan in the Recombination Detection Program (RDP) v.4.87. Parameters for bootscan analysis were: window size = 200; step size = 20; bootstrap replicates = 1,000; cutoff percentage = 70; use neighbor joining trees; calculate binomial p-value; model option = Kimura 1980 [21]. For each analysis, a single alignment was created which included the modern haplogroup, all 9 Neanderthal and all 6 Ancient *H. sapiens* sequences. When rCRS was used as query, two sets of possible parental sequences were selected: either Neanderthals Mezmaiskaya and Altai and ancient *H. sapiens* Fumane and Ust Ishim or only Neanderthals Feldhofer1, Mezmaiskaya and Vindija 33.16. For haplogroups L0d1a and L3d3b possible parental sequences were Neanderthals Feldhofer1, Mezmaiskaya and Vindija 33.16 and Ancient *H. sapiens* Kostenki 14, Fumane, Doni Vestonice 14 and Tianyuan. For haplogroups M29a and R0a possible parental sequences were Neanderthals Mezmaiskaya and Altai and Ancient *H. sapiens* Kostenki 14 and Doni Vestonice 14. For haplogroup N1b1a3 possible parental sequences were Neanderthals Feldhofer1 and Vindija 33.16 and Ancient *H. sapiens* Kostenki 14 and Doni Vestonice 14.

### 2.5. Statistical analysis

For variants (SNVs) calling three different datasets were used: (1) the whole mitogenome from the 102 sequences alignment; (2) the 128 to 315bp fragment and (3) the 6,950 to 7,660bp fragment in the same alignment. All fasta alignments were processed using the MSA2VCF software to generate the VCF files [22]. The options used on msa2vcf were: --haploid --output. To convert the VCF files to Plink format we used the vcftools package [23]. The whole mitogenome alignment with 103 sequences had 785 SNPs (positions containing gaps in at least one sequence were excluded from the analysis). Both 128 – 315 and 6,950-7,660 fragments had 24 SNPs.

Principal component analysis (PCA) was performed using the PLINK software v1.90b4 [24,25]. PCA figure plotting was made using Genesis PCA and admixture plot viewer (http://www.bioinf.wits.ac.za/software/genesis/). The first two principal components were chosen for the Neanderthal - *H. sapiens* comparison.

## 3. Results

### 3.1. Distribution of Neanderthal SNVs

An alignment of 102 sequences was produced which contained the mitogenomes of 41 archaic AMH (Table 1), 9 *Homo sapiens neanderthalensis* (Table 1) and 52 contemporary *Homo sapiens sapiens* with representatives of the major worldwide mitochondrial haplogroups (Table 2). This alignment contained the revised Cambridge Reference Sequence (rCRS) used as numbering reference for all polymorphisms identified [16]. We found 66 positions in which contemporary *Homo sapiens sapiens* were identical with at least one *Homo sapiens neanderthalensis* sequence position (Figure 1, Tables 3 and 4). In 13 positions just a subset of contemporary *Homo sapiens sapiens* were identical with at least one *Homo sapiens neanderthalensis* sequence position and at least one sequence of archaic AMH. In 175 positions *Homo sapiens neanderthalensis* differed from contemporary *Homo sapiens sapiens* and archaic AMH. In 11 positions the archaic AMH differed from other sequences and in 653 positions the contemporary *Homo sapiens sapiens* presented expected variants among haplogroups that were not relevant for this analysis.

**Fig. 1.**
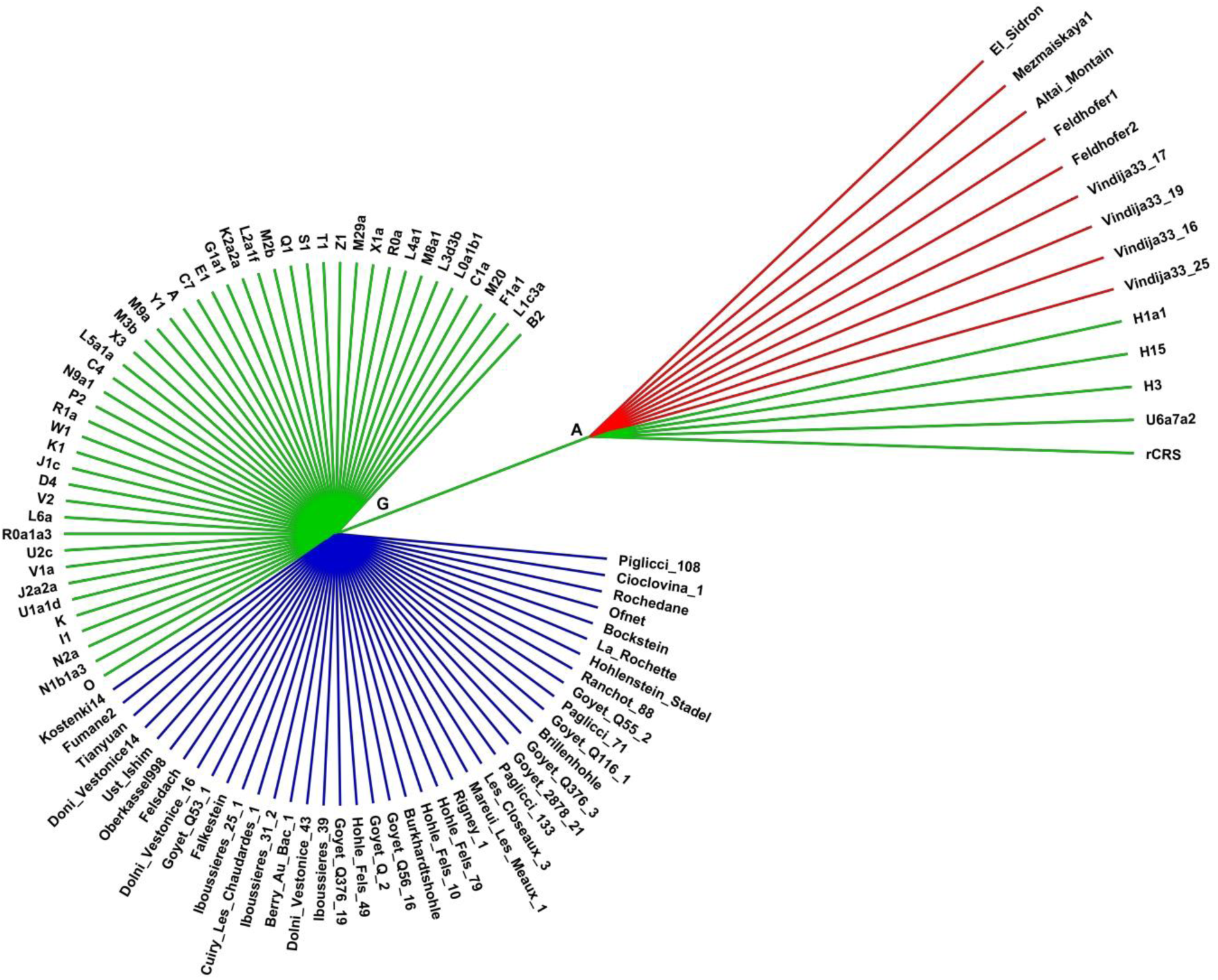
Cladogram of mitochondrial genome position 2,706. This N-SNV is a G-A transition in the 16S rRNA gene. This genetic signature (Neanderthal signature 2706G) is present in all Neanderthal sequences (Red branches) and in European haplogroups U (<u>U</u>5a7a2) and H (<u>H</u>1a1, <u>H</u>3, <u>H</u>15), including the Revised Cambridge Reference Sequence (rCRS, haplogroup <u>H</u>2a2a1). The Neanderthal signature 2706G is absent in all archaic AMH (Anatomically Modern Humans from Europe in temporal overlap with Neanderthals - Blue branches) and other modern mitochondrial haplogroups. Position numbering corresponds to rCRS positions. This cladogram is representative of the 66 cladograms generated for each N-SNV giving similar topologies and which are available upon request.

**Table 3.**
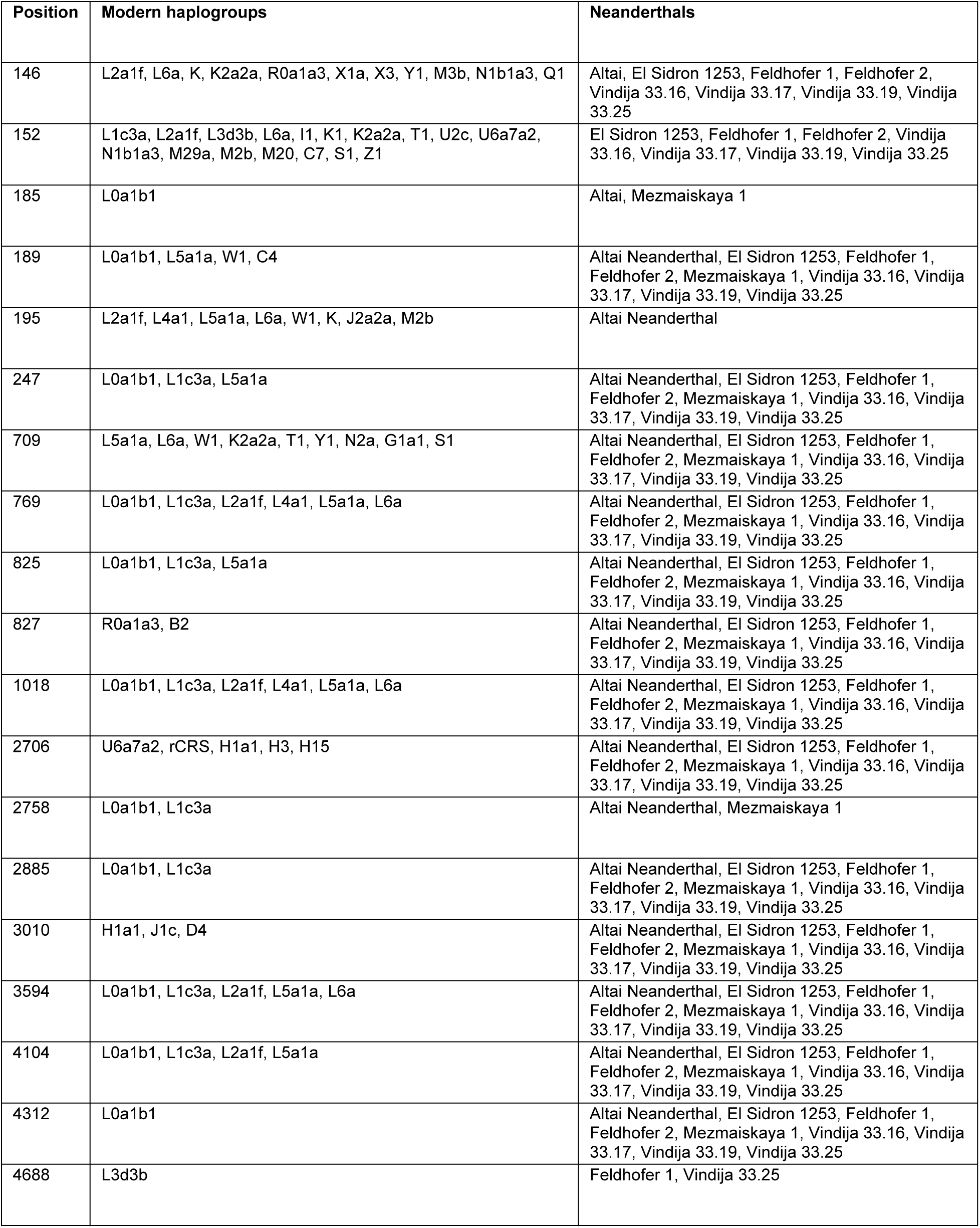
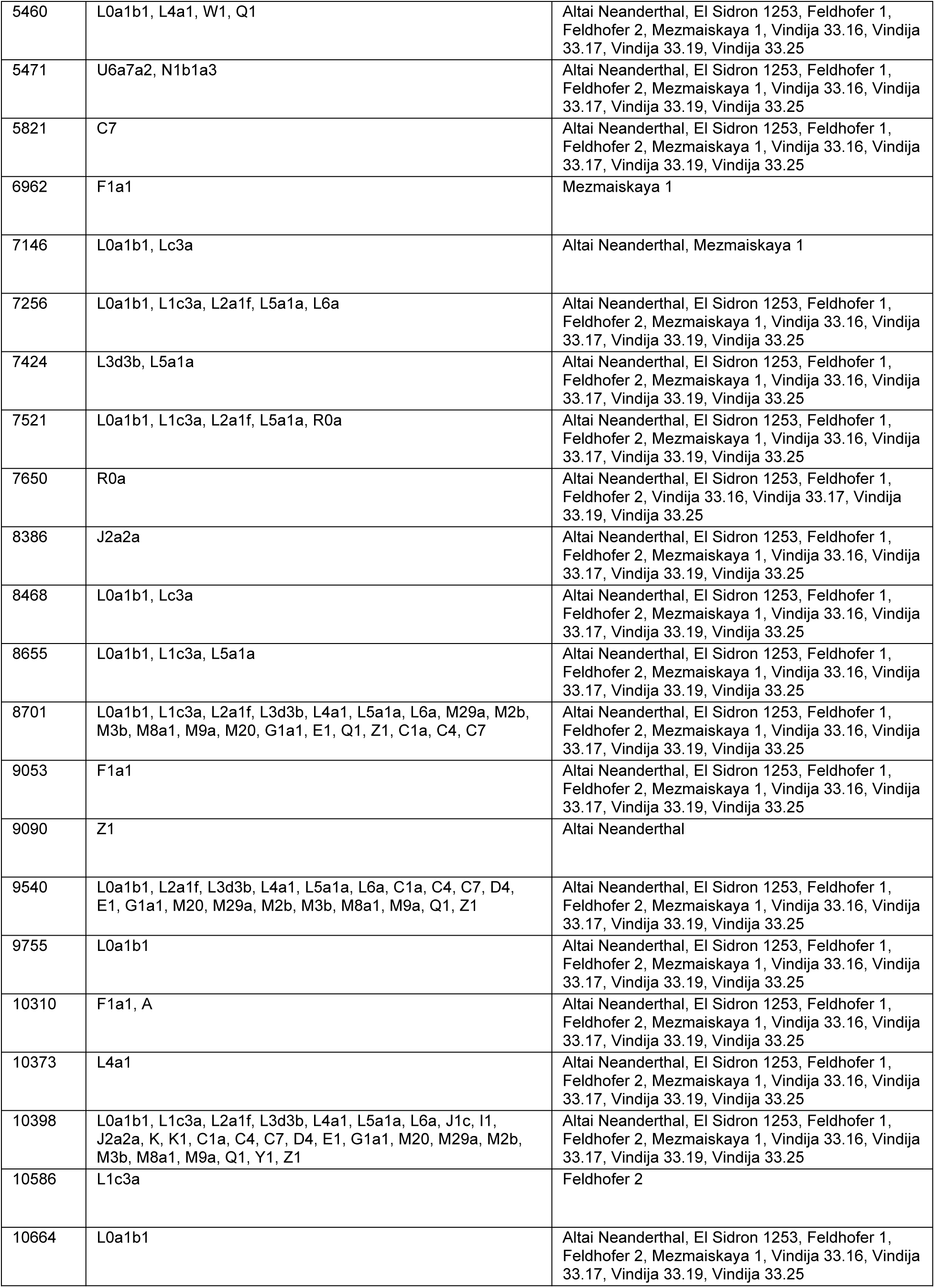
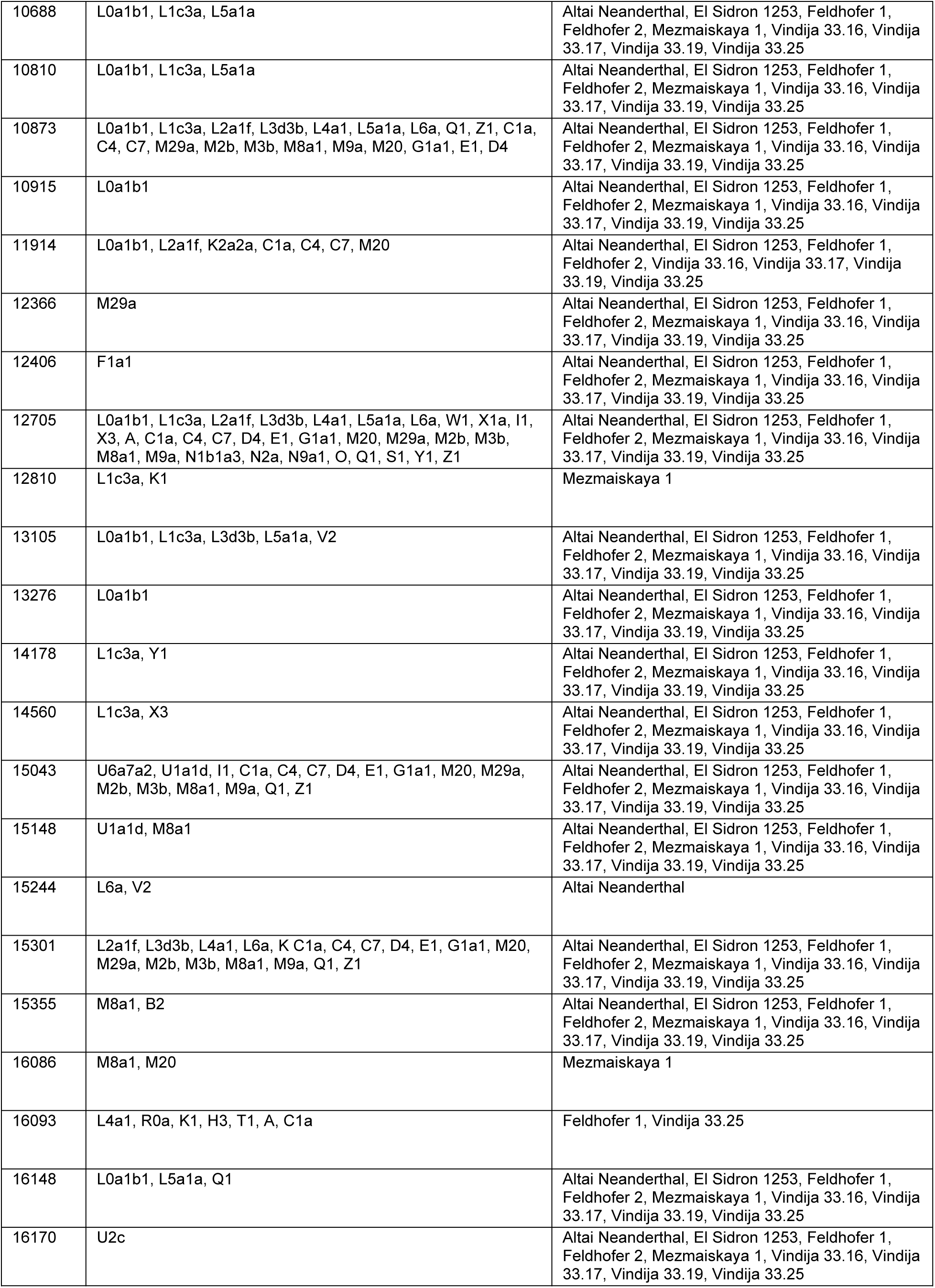
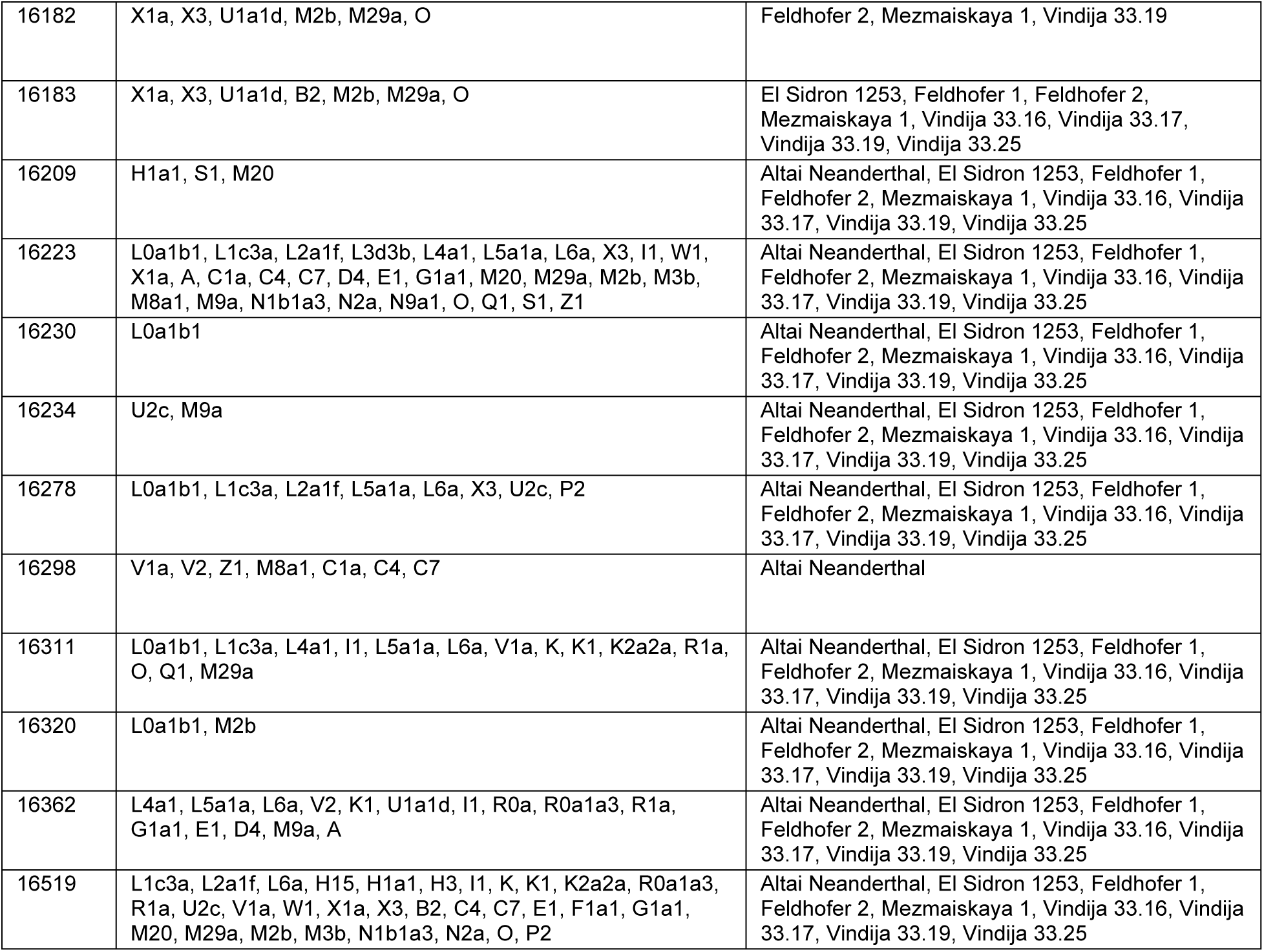
Complete list of mitogenome positions where present day sequences are identical with Neanderthals.

**Table 4.** Full list of all mitogenome N-SNVs analysed in this study. Ts=transition, Tv=transversion, AA=amino acid, Nt=nucleotide, Freq=Frequency, Alig.=alignment, 1K Gen.=1000 genomes project database, Prot. Effect=Protein effect.

| Position | DbSNP | Change | Change | Change | Type | Prot. Effect | Alignment. | 1K Gen. | Gene | Codon |
| --- | --- | --- | --- | --- | --- | --- | --- | --- | --- | --- |
|  | rs | AA | Nt | Codon |  |  | Freq (%) | Freq (%) |  | Position |
| 146 | rs370482130 | - | T>C | - | Ts | - | 28.40% | 21.40% | - | - |
| 150 | rs62581312 | - | C > T | - | Ts | - | 20.90% | 19.10% | - | - |
| 152 | rs117135796 | - | T>C | - | Ts | - | 38.80% | 33% | - | - |
| 185 | rs879015046 | - | G>A | - | Ts | - | 4.50% | 3.70% | - | - |
| 189 | rs371543232 | - | A>G | - | Ts | - | 19.40% | 7.70% | - | - |
| 195 | rs2857291 | - | T>C | - | Ts | - | 13.40% | 24.70% | - | - |
| 247 | rs41334645 | - | G>A | - | Ts | - | 17.90% | 7.00% | - | - |
| 709 | rs2853517 | - | G>A | - | Ts | - | 26.90% | 16.70% | - | - |
| 769 | rs2853519 | - | G>A | - | Ts | - | 22.40% | 16.40% | - | - |
| 825 | rs2853520 | - | T>A | - | Tv | - | 17.90% | 7.10% | - | - |
| 827 | rs28358569 | - | A>G | - | Ts | - | 16.40% | 4.30% | - | - |
| 1018 | rs2856982 | - | G>A | - | Ts | - | 22.40% | 16.50% | - | - |
| 2706 | rs2854128 | - | A>G | - | Ts | - | 80.60% | 89.00% | - | - |
| 2758 | rs2856980 | - | G>A | - | Ts | - | 6.00% | 6.60% | - | - |
| 2885 | rs2854130 | - | T>C | - | Ts | - | 16.40% | 6.80% | - | - |
| 3010 | rs3928306 | - | G>A | - | Ts | - | 17.90% | 11.20% | - | - |
| 3594 | rs193303025 | - | C>T | GTC>GTT | Ts | No change | 20.90% | 15.90% | ND1 | 3 |
| 4104 | rs1117205 | - | A>G | CTA>CTG | Ts | No change | 19.40% | 15.80% | ND1 | 3 |
| 4312 | rs193303033 | - | C>T | - | Ts | - | 14.90% | 1.80% | tRNA | - |
| 4688 | rs878853056 | - | T>C | GCT>GCC | Ts | No change | 4.50% | 0.40% | ND2 | 3 |
| 5460 | rs3021088 | A>T | G>A | GCC>ACC | Ts | AA change | 19.40% | 7.50% | ND2 | 1 |
| 5471 | rs879108598 | - | G>A | ACG>ACA | Ts | No change | 16.40% | 0.50% | ND2 | 3 |
| 5821 | rs56133209 | - | G>A | - | Ts | - | 14.90% | 0.50% | tRNA | - |
| 6962 | rs1970771 | - | G>A | CTG>CTA | Ts | No change | 3.00% | 3.00% | COX1 | 3 |
| 7146 | rs372136420 | T>A | A>G | ACT>GCT | Ts | AA change | 6.00% | 6.10% | COX1 | 1 |
| 7256 | - | - | C>T | AAC>AAT | Ts | No change | 20.90% | 15.80% | COX1 | 3 |
| 7424 | - | - | A>G | GAA>GAG | Ts | No change | 16.40% | 2.30% | COX1 | 3 |
| 7521 | rs199474817 | - | G>A | - | Ts | - | 20.90% | 16.00% | tRNA | - |
| 7650 | - | T>I | C>T | ACC>ATC | Ts | AA change | 13.40% | 0.00% | COX2 | 2 |
| 8386 | - | - | C>T | ACC>ACT | Ts | No change | 14.90% | 0.00% | ATP8 | 3 |
| 8468 | rs1116907 | - | C>T | CTA>TTA | Ts | No change | 16.40% | 6.70% | ATP8 | 1 |
| 8655 | - | - | C>T | ATC>ATT | Ts | No change | 17.90% | 7.10% | ATP6 | 3 |
| 9053 | rs199646902 | S>N | G>A | AGC>AAC | Ts | AA change | 14.90% | 2.60% | ATP6 | 2 |
| 9090 | rs386829064 | - | T>C | TCT>TCC | Ts | No change | 3.00% | 0.30% | ATP6 | 3 |
| 9755 | rs2856985 | - | G>A | GAG>GAA | Ts | No change | 14.90% | 2.00% | COX3 | 3 |
| 10310 | rs41467651 | - | G>A | CTG>CTA | Ts | No change | 16.40% | 4.00% | ND3 | 3 |
| 10373 | rs28358277 | - | G>A | GAG>GAA | Ts | No change | 14.90% | 3.00% | ND3 | 3 |
| 10586 | rs28358281 | - | G>A | TCG>TCA | Ts | No change | 3.00% | 2.60% | ND4L | 3 |
| 10664 | - | - | C>T | GTC>GTT | Ts | No change | 14.90% | 1.80% | ND4L | 3 |
| 10688 | rs2853488 | - | G>A | GTG>GTA | Ts | No change | 17.90% | 7.00% | ND4L | 3 |
| 10810 | rs28358282 | - | T>C | CTT>CTC | Ts | No change | 17.9 | 7.10% | ND4 | 3 |
| 10915 | rs2857285 | - | T>C | TGT>TGC | Ts | No change | 14.90% | 2.30% | ND4 | 3 |
| 11914 | rs2853496 | - | G>A | ACG>ACA | Ts | No change | 23.90% | 12.70% | ND4 | 3 |
| 12366 | - | - | A>G | CTA>CTG | Ts | No change | 14.90% | 0.20% | ND5 | 3 |
| 12810 | rs28359174 | - | A>G | TGA>TGG | Ts | No change | 4.50% | 2.40% | ND5 | 3 |
| 13105 | rs2853501 | I>V | A>G | ATC>GTC | Ts | AA change | 20.90% | 13.70% | ND5 | 1 |
| 13276 | rs2853502 | M>V | A>G | ATA>GTA | Ts | AA change | 14.90% | 1.80% | ND5 | 1 |
| 14178 | rs28357671 | I>V | T>C | ATT>GTT | Ts | AA change | 16.40% | 5.20% | ND6 | 1 |
| 14560 | rs28357676 | - | G>A | GTC>GTT | Ts | No change | 16.40% | 5.40% | ND6 | 3 |
| 15148 | rs527236206 | - | G>A | CCG>CCA | Ts | No change | 16.40% | 1.10% | CYTB | 3 |
| 15244 | rs28357369 | - | A>G | GGA>GGG | Ts | No change | 4.50% | 2.10% | CYTB | 3 |
| 15355 | rs527236181 | - | G>A | ACG>ACA | Ts | No change | 16.40% | 0.60% | CYTB | 3 |
| 16086 | rs386420030 | - | T>C | - | Ts | - | 4.50% | 2.10% | - | - |
| 16093 | rs2853511 | - | T>C | - | Ts | - | 13.40% | 5.60% | - | - |
| 16148 | rs201893071 | - | C>T | - | Ts | - | 17.90% | 3.10% | - | - |
| 16169 | rs879121566 | - | C>T | - | Ts | - | 1.50% | 0.80% | - | - |
| 16182 | rs879044186 | - | A>C | - | Tv | - | 15.20% | 2.70% | - | - |
| 16183 | rs28671493 | - | A>G | - | Ts | - | 1.50% | 0.40% | - | - |
| 16209 | rs386829278 | - | T>C | - | Ts | - | 17.90% | 3.00% | - | - |
| 16230 | rs2853514 | - | A>G | - | Ts | - | 14.90% | 2.10% | - | - |
| 16234 | rs368259300 | - | T>C | - | Ts | - | 16.40% | 1.90% | - | - |
| 16278 | rs41458645 | - | C>T | - | Ts | - | 25.40% | 20.80% | - | - |
| 16298 | rs148377232 | - | T>C | - | Ts | - | 11.90% | 5.00% | - | - |
| 16311 | rs34799580 | - | T>C | - | Ts | - | 34.30% | 18.70% | - | - |
| 16320 | rs62581338 | - | C>T | - | Ts | - | 16.40% | 5.20% | - | - |
| 16519 | rs3937033 | - | T>C | - | Ts | - | 61.20% | 62.30% | - | - |

To depict the distribution of Neanderthal SNVs, or N-SNVs, in different human mitochondrial haplogroups, we constructed a heat map of N-SNVs (Figure 2). Most N-SNVs are concentrated in the D-loop, followed by 12SrDNA and 16SrDNA. Among tRNAs N-SNVs were found only in isoleucine, asparagine and cysteine tRNAs. In *COX2* only one N-SNV was found in haplogroup R0a.

**Fig. 2.**
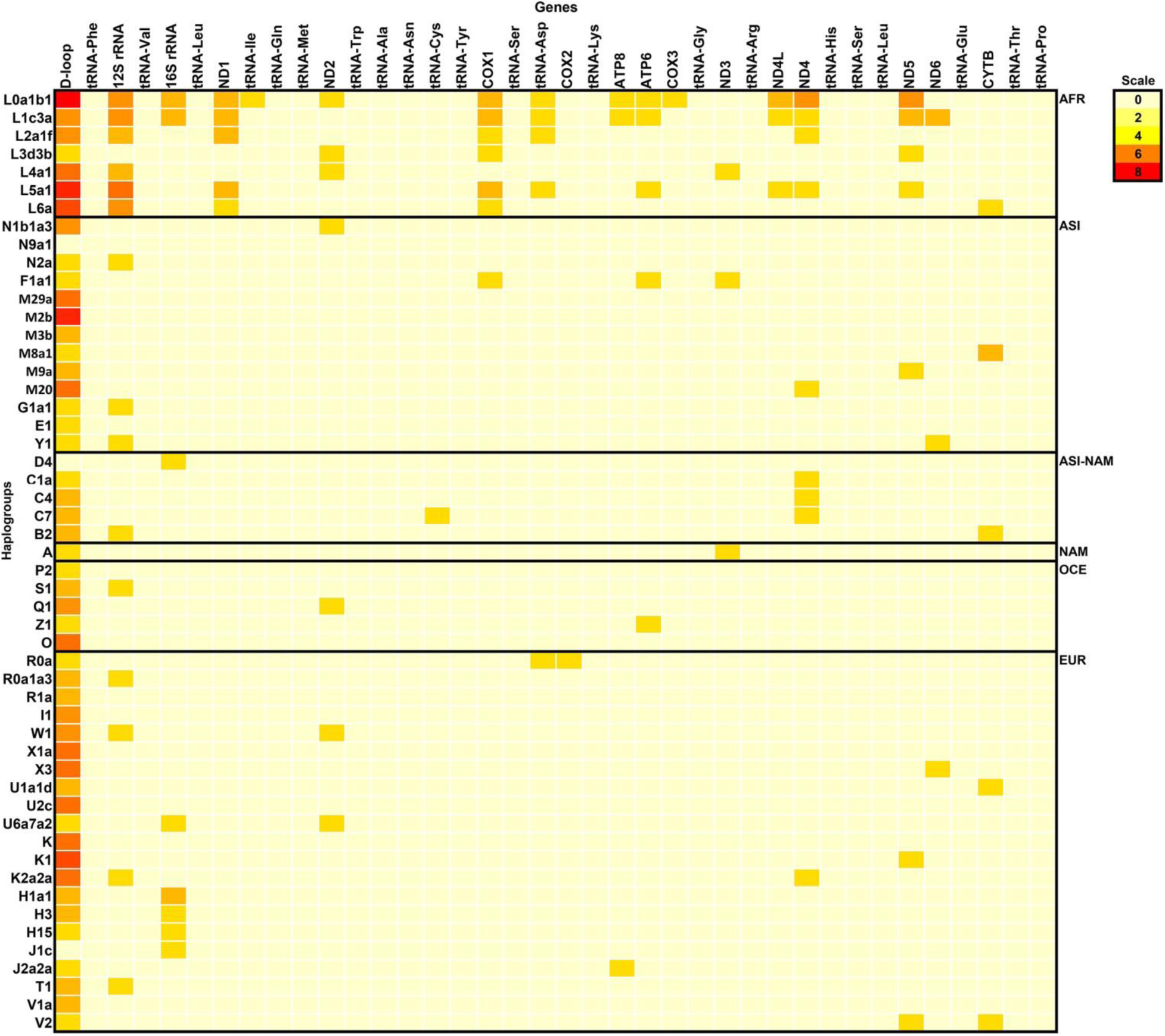
Heatmap of Neanderthal signatures along the mitochondrial genome in different haplogroups. The color scale indicates the number of Neanderthal signatures (N-SNVs) present in modern human mitochondrial haplogroups. These signatures are absent in Ancient *H. sapiens* whose time range overlapped with Neanderthals in Europe (from approximately 50,000 to 28,000 years ago, Table 1). AFR=African, ASI=Asian, MDE=Middle East, NAM=Native American, OCE=Oceania and EUR=European.

Of the 66 N-SNVs identified, 20 are common to modern African and Eurasian haplogroups, 25 are exclusive to African haplogroups and 21 are exclusive to Eurasian haplogroups. In Figure 3 the distribution of N-SNVs is depicted in the human mitogenome map. The distribution reveals that 11 Eurasian N-SNVs are in coding regions, 3 in rRNA genes and 1 in a tRNA gene.

**Fig. 3.**
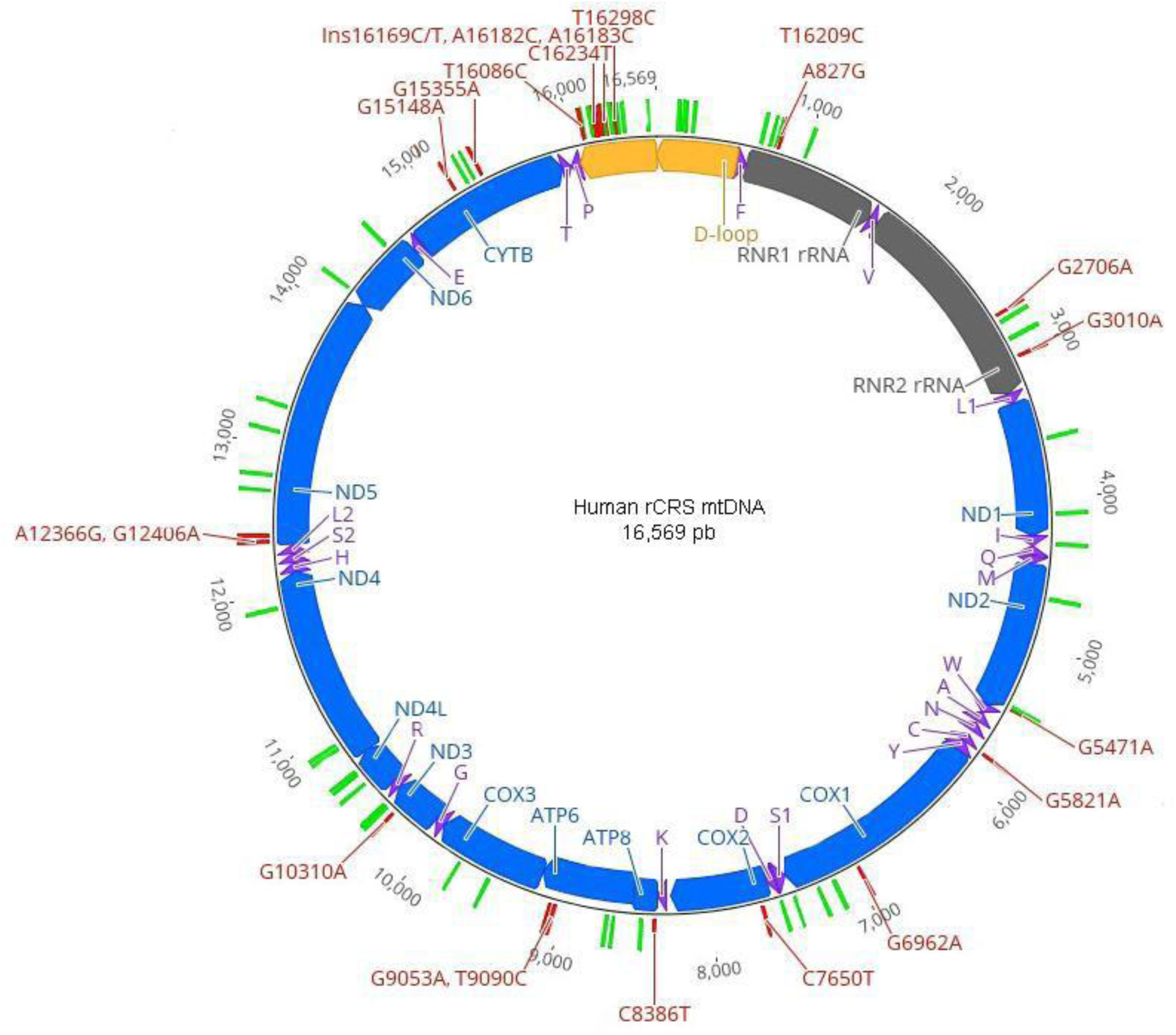
Position of Neanderthal SNVs (N-SNVs) in the mitochondrial genome. Blue bars indicate protein coding genes, gray bars indicate ribosomal RNA genes, purple arrows indicate tRNAs, yellow bars indicate the D-loop, green bars indicate Neanderthal SNVs present in African haplogroups and red bars indicate N-SNVs exclusive of Eurasian haplogroups.

The distribution of N-SNVs in modern haplogroups and archaic AMH can be summarized as possible six introgression routes with predicted consequences (Figure 4). In hypothesis (1) SNVs conserved between Neanderthals, archaic AMH and present-day humans should be observed in all sequences; in (2) if the introgression crosses occurred between Neanderthals and archaic AMH prior to divergence that separated archaic AMH from lineages of African and Eurasian haplogroups it is expected that N-SNVs would be present in all sequences, as observed for 13 N-SNVs. In (3) the crosses would have occurred only with archaic AMH who did not contribute to present day haplogroups while hypotheses (4), (5) and (6) would represent crosses between Neanderthals and lineages who contributed to present day haplogroups. The 66 N-SNVs are consistent with these hypotheses, which exclude their presence in archaic AMH and thus are likely signals of crosses between these two subspecies of humans. More importantly no SNVs are identical between Neanderthals and archaic AMH at the exclusion of present day mitogenomes of either African and Eurasian haplogroups which suggests that the presence of N-SNVs must be a signal of horizontal transfer or a very high number of reverse/convergent substitutions.

**Fig. 4.**
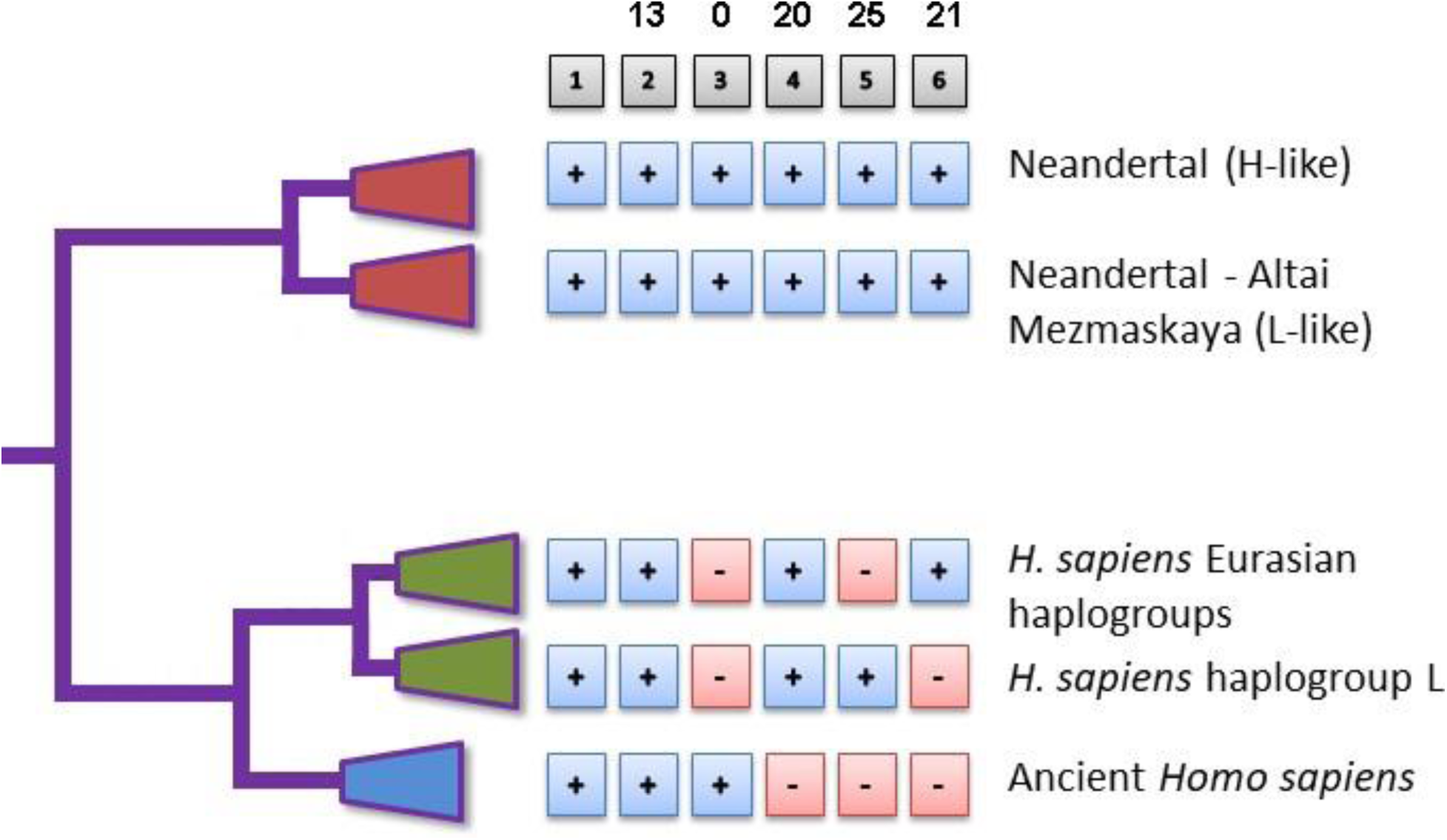
Phylogenetic distribution of Neanderthal variants in modern humans and archaic AMH. The diagram depicts six possible introgression routes, or intercross events, between Neanderthals, archaic AMH and lineages leading to present day *Homo sapiens* haplogroups. Numbers above gray boxes indicate the number of specific N-SNVs in that particular clade. Plus (+) and minus (-) signs within blue and red boxes indicate N-SNVs presence or absence in that particular clade.

### 3.2. Disease associated N-SNVs

Among the 66 N-SNVs, 7 are associated with diseases as depicted in Table 5. Of note, 4 of these disease-associated N-SNVs were observed in African haplogroups (L0, L1, L2, L3, L4, L5 and L6) and 3 were observed exclusively in Eurasian haplogroups. The most common diseases associated with N-SNVs are neurological disorders and tumors. One N-SNVs, in position 15,043 and associated with depression, was also found in one archaic AMH. Although not considered a *bona fide* N-SNV based on our exclusion criteria, it is relevant because chronic depression has been associated with Neanderthal introgression [26]. Among protein coding genes an important N-SNV was found in *ND2* (position 5,460) causing an amino acid change from alanine to threonine that has been associated with Alzheimer’s disease and Parkinson’s disease due to its high prevalence in the brains of Alzheimer’s and Parkinson’s patients [27–29]. This amino acid substitution changes from a nonpolar amino acid to a polar amino acid, which promotes destabilizing effects in the encoded NADH dehydrogenase. Nonpolar to polar amino acid changes are associated with amyloid diseases [30–32]. Other disease associated N-SNVs are in the D-loop, 16S rRNA and tRNA-Cys (Table 2). The prevalence of diseases associated with N-SNVs in Table 2 are (data from https://vizhub.healthdata.org/gbd-compare/, accessed on August 31 2019): Bipolar disorder = 596 cases in 100,000 persons (596/100,000), Parkinsons’ disease = 111/100,000, Alzheimer’s disease = 588/100,000, Melanoma = 30.42/100,000, Ovarian cancer = 17.71/100,000, Lung cancer = 43/100,000, Prostate cancer = 129/100,000, Stomach cancer = 36.9/100,000. Also, Cycling vomiting syndrome 3.2/100,000 [33], Deafness = 110/1,000 [34], Glioblastoma = 10 /100,000 [35]

**Table 5.** Disease associated Neanderthal SNVs. Associations as compiled and summarized in MITOMAP. [18].

|  |  |  |  |  |  |  |  |
| --- | --- | --- | --- | --- | --- | --- | --- |
| Position | 195 | 3,010 | 5,460 | 5,821 | 16,093 | 16,183 | 16,519 |
| Region / gene | D-loop | 16S rRNA | ND2 | tRNA-Cys | D-loop | D-loop | D-loop |
| dbSNP rs | rs2857291 | rs3928306 | rs3021088 | rs56133209 | rs2853511 | rs28671493 | rs3937033 |
| DNA change | T>C | G>A | G>A | G>A | T>C | A>C | T>C |
| Type | Transition | Transition | Transition | Transition | Transition | Transversion | Transition |
| Codon position | - | - | 1 | - | - | - | - |
| Codon effect | - | - | Non Syn. | - | - | - | - |
| Codon change | - | - | GCC>ACC | - | - | - | - |
| Protein change | - | - | Ala>Thr | - | - | - | - |
| Disease associations | Bipolar disorder / melanoma patients | Cyclic Vomiting Syndrome with Migraine | Alzheimer's disease / Parkinson's disease | Deafness helper mutation | Cyclic Vomiting Syndrome | Melanoma patients | Cyclic Vomiting Syndrome with Migraine /metastasis / glioblastoma, gastric, lung, ovarian, prostate tumors |
| Modern haplogroups | L2a1f, L4a1, L5a1a, L6a, W1, K, J2a2a, M2b | H1a1, J1c, D4 | L0a1b1, L4a1, W1, Q1 | C7 | L4a1, R0a, K1, H3, T1, A, C1a | X1a, X3, U1a1d, B2, M2b, M29a, O | L1c3a, L2a1f, L6a, H15, H1a1, H3, I1, K, K1, K2a2a, R0a1a3, R1a, U2c, V1a, W1, X1a, X3, B2, C4, C7, E1, F1a1, G1a1, M20, M29a, M2b, M3b, N1b1a3, N2a, O, P2 |
| Neanderthal | Altai | Altai<br>El Sidron 1253<br>Feldhofer 1<br>Feldhofer 2<br>Mezmaiskaya1<br>Vindija 33.16<br>Vindija 33.17<br>Vindija 33.19<br>Vindija 33.25 | Altai<br>El Sidron 1253<br>Feldhofer 1<br>Feldhofer 2<br>Mezmaiskaya1<br>Vindija 33.16<br>Vindija 33.17<br>Vindija 33.19<br>Vindija 33.25 | Altai<br>El Sidron 1253<br>Feldhofer 1<br>Feldhofer 2<br>Mezmaiskaya1<br>Vindija 33.16<br>Vindija 33.17<br>Vindija 33.19<br>Vindija 33.25 | Feldhofer1<br>Vin. 33.25 | El Sidron 1253<br>Feldhofer 1<br>Feldhofer 2<br>Mezmaiskaya1<br>Vindija 33.16<br>Vindija 33.17<br>Vindija 33.19<br>Vindija 33.25 | Altai<br>El Sidron 1253<br>Feldhofer 1<br>Feldhofer 2<br>Mezmaiskaya1<br>Vindija 33.16<br>Vindija 33.17<br>Vindija 33.19<br>Vindija 33.25 |

### 3.3. Haplogroups of paleogenomes

Mitogenomes of Neanderthals and archaic AMH were classified in haplogroups according to sequence similarity with extant human mitogenomes using Haplogrep 2 [20] (Table 1). 83% of archaic AMH mitogenomes belong to haplogroup U (45% haplogroup U5 and 16% to haplogroup U2) which is consistent as U being the oldest European haplogroup. Among the 9 Neanderthal mitogenomes, the 7 more recent genomes can be classified as haplogroup H1 (European) while the two oldest can be classified as haplogroup L (African) (Table 1). N-SNVs at positions 16,278 and 16,298 are associated with intelligence quotient [36]. N-SNV 16,278 is found in African haplogroups (L0, L1, L2, L5 and L6) and two Eurasian haplogroups (X3, U2c and P2) and in all Neanderthal sequences while N-SNV 16.298 is found only in Eurasian-Native American haplogroups (V1, V2, M8, C1, C4, C7 and Z1) and only in the Altai Neanderthal.

### 3.4. PCAs of Neanderthal and Human mitogenomes

Principal component analysis (PCA) of the whole mitochondrial genome shows four clusters: (1) the modern haplogroups including the ancient *H. sapiens* (Table 1), (2) the L haplogroup cluster, (3) the Neanderthal Altai-Mezmaskaya L-like cluster (Table 1) and (4) the Neanderthal H-like group (Figure 5). However, the PCA of the segment corresponding to the ribosomal RNA gene proximal half produces a pattern that approximates the L haplogroup cluster to the Neanderthal Altai-Mezmaskaya L-like cluster suggesting the introgression point. The PCA of ribosomal RNA gene distal half suggests an opposite pattern with L cluster closer to Neanderthal although not as close as shown in Figure 5B.

**Fig. 5.**
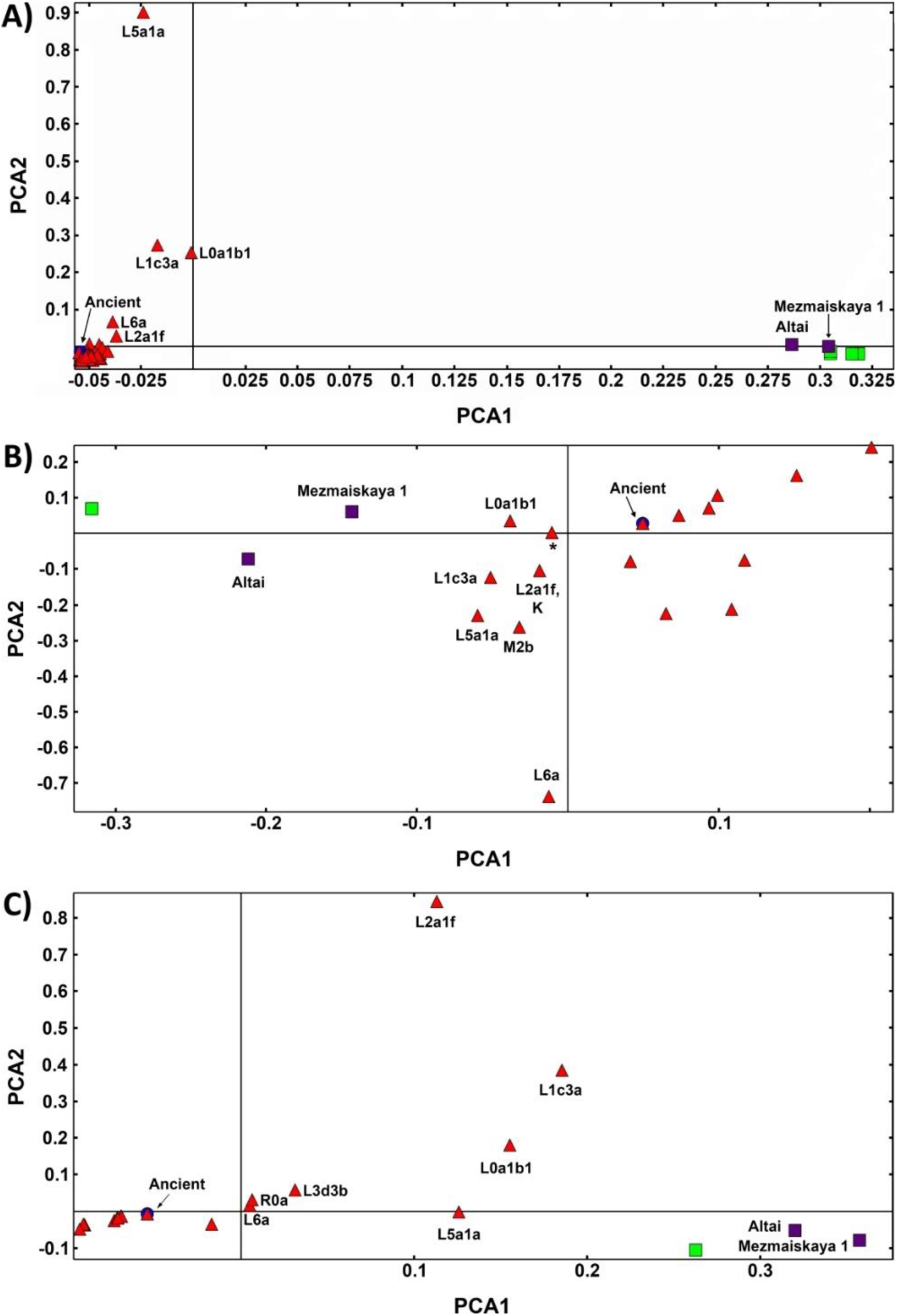
Principal Component Analysis (PCA) of Human and Neanderthal mitochondrial genomes. In (A) PCA results for all 16,569 positions of 103 mitogenomes being 9 Neanderthals (green squares for H-like sequences and purple squares for L-like sequences), 42 archaic AMH (blue circles) and 53 modern *H. sapiens* haplogroups (red triangles). X-Axis denotes the value for PC1, while y-Axis denotes values for PC2. Each dot in the figure represents one or more individuals. In (B) PCA results for the segment comprising positions 128 to 315 extracted from the 103 mitogenomes alignment and in (C) PCA results for the segment comprising positions 6,950 to 7,660 extracted from the 103 mitogenomes alignment.

### 3.5. Bootscan analysis of mitogenomes

We tested potential recombination in our dataset with bootscan. We used a different set of parental sequences depending on the query mitogenome (Figure 6). The bootscan analysis indicates that there are potential recombination points. Upon deeper analysis we observed that bootscan considers the Neanderthal specific signatures, such as in L haplogroups, as recombination points. Although the bootscan putative recombination segments are above the bootstrap threshold we do not consider this as definitive evidence of recombination since the segments between the Neanderthal signatures are almost identical. Bootscan analysis excluded Human-Neanderthal recombination in rCRS sequence (Figure 7). The sensivity of bootscan to substitution models and alignment methods was assessed by comparing the same set query-parentals with different parameters (Figure 8), revealing minor profile alterations. The alignment parameters are not so critical in this case because the sequences are extremely conserved (918 polymorphic positions in 16,565bp). Although indels are present in the alignments, 99% are located near the H promoter in the D-loop region. These are automatically excluded in phylogeny inference algorithms and therefore have no weight in bootscan results. The “positional homology” is therefore solid, particularly in coding domains and regions without repeats in non-coding domains. The Neanderthal signatures are in unambiguously aligned segments.

**Fig. 6.**
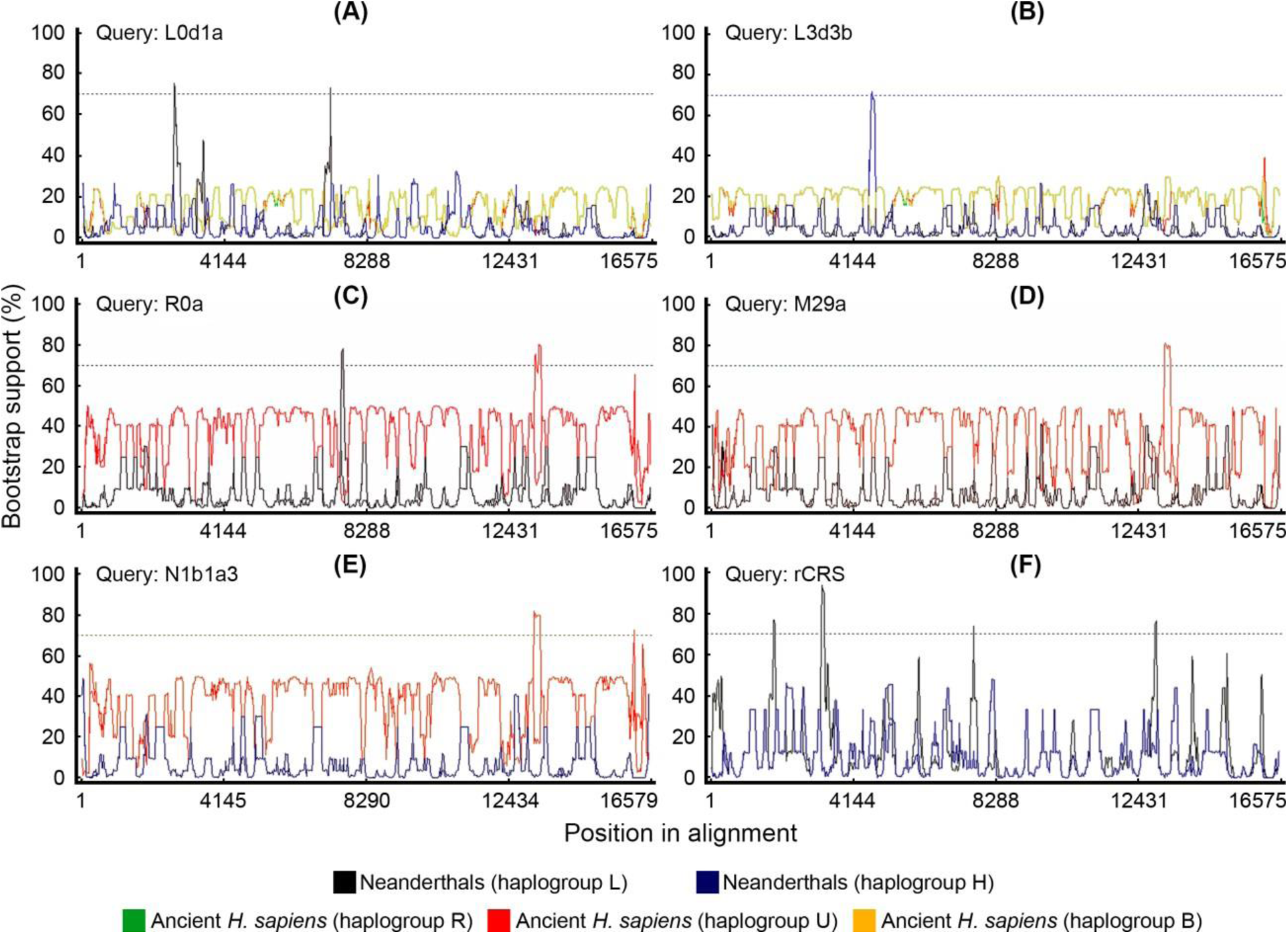
Bootscan recombination analysis. Possible recombination points in were detected using different modern haplogroups as queries and Ancient *H. sapiens* or Neanderthals as putative parental sequences.

**Fig. 7.**
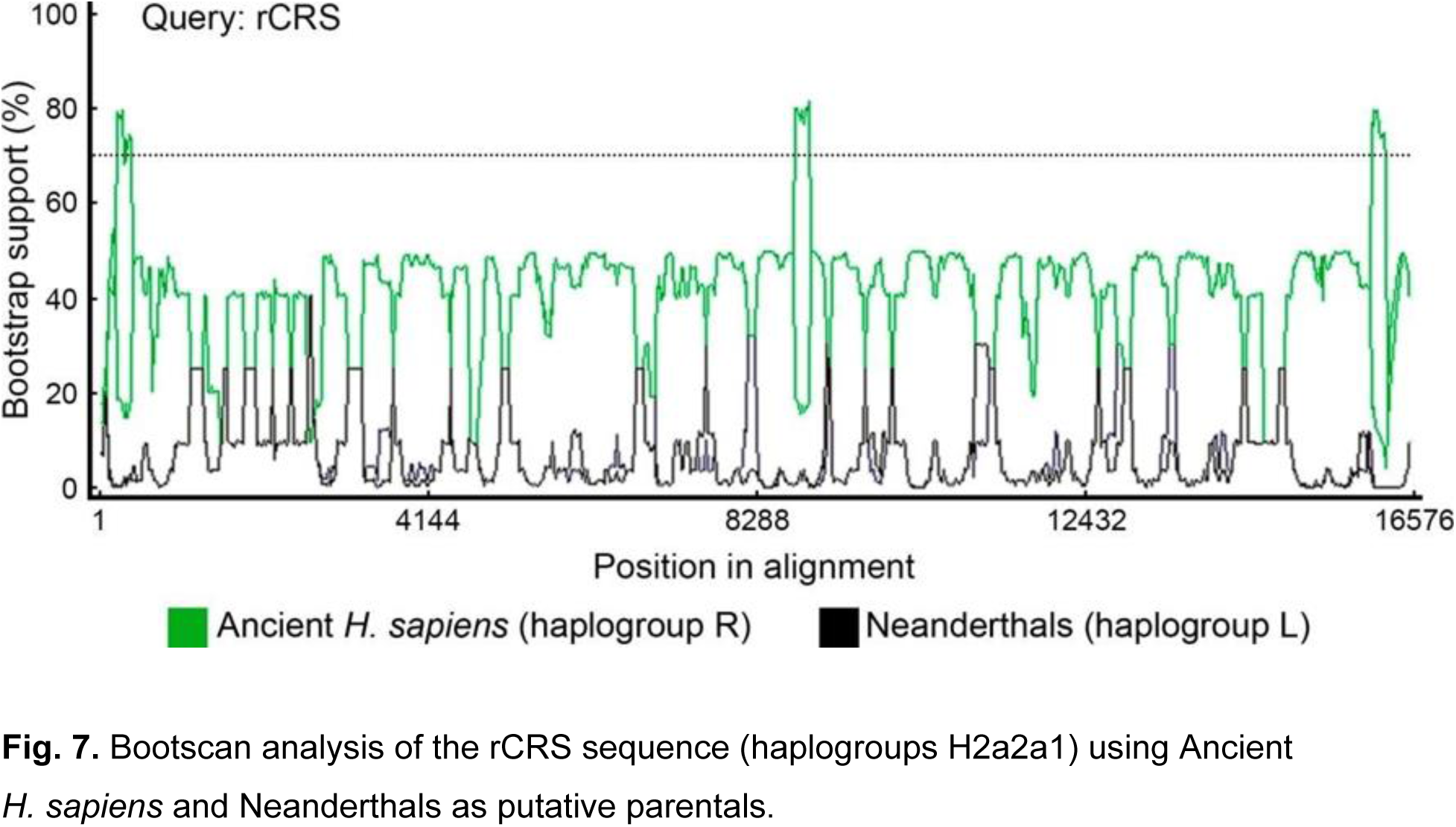
Bootscan analysis of the rCRS sequence (haplogroups H2a2a1) using Ancient *H. sapiens* and Neanderthals as putative parentals.

**Fig. 8.**
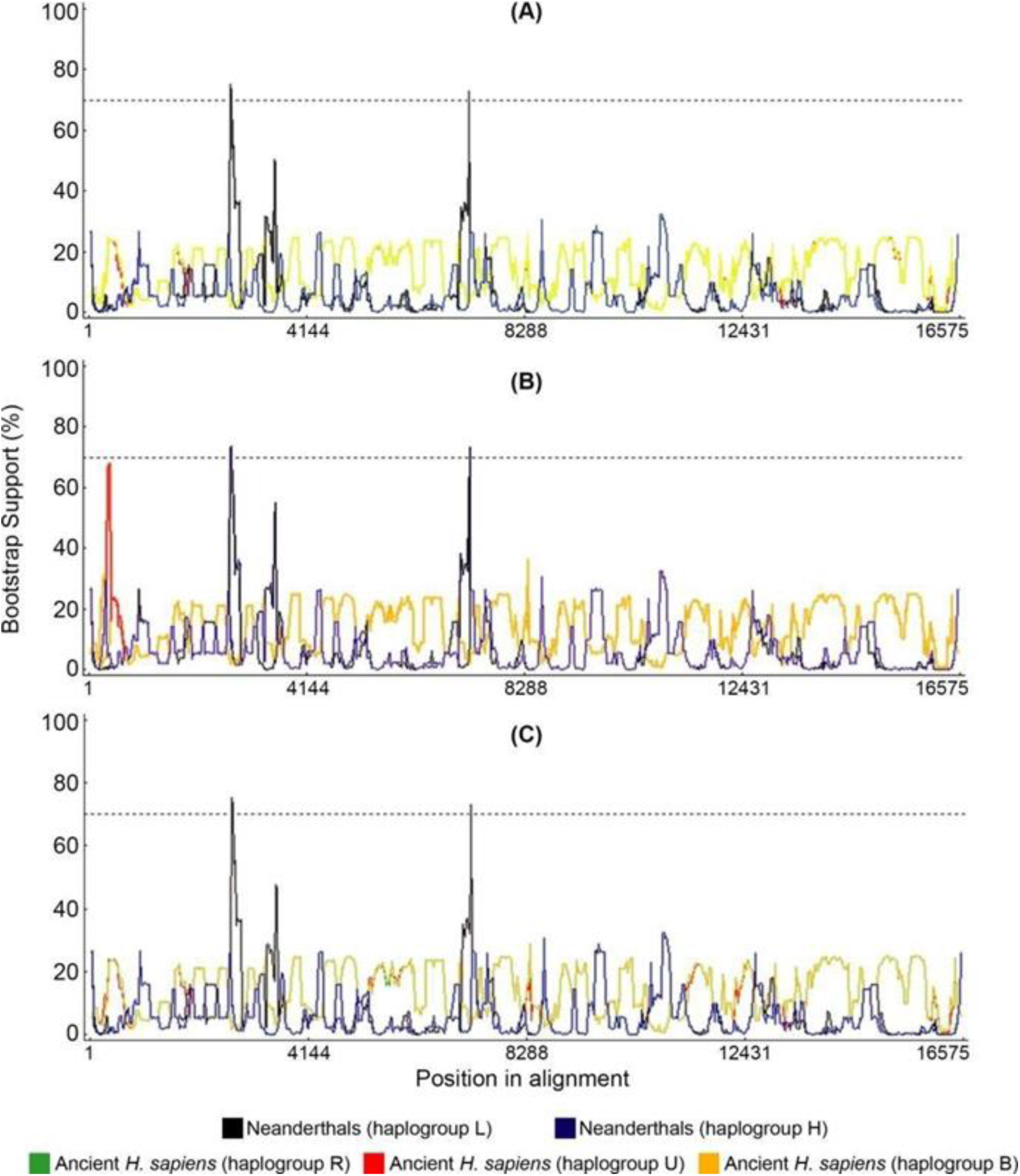
Bootscan/RDP analysis of haplogroup L0d1a showing the profile consistency regardless the alignment algorithm or bootscan model option. (A) Map to reference aligment and Felsenstein model on bootscan; (B) MAFFT alignment with Kimura two parameters model on bootscan; (C) Map to reference aligment and Kimura two parameters model on bootscan. Neanderthal sequences from haplogroup L (black line) and haplogroup H (blue line) and Ancient *H. sapiens* from haplogroup R (green line), haplogroup U (red line) and haplogroup B (orange line) were used as parentals. Neanderthal sequences were Feldhofer 1, Mezmaiskaya and Vindija33_16 and ancient *H. sapiens* sequences were Kostenki, Fumane, Doni Vestonice 14 and Tianyuan.

## 4. Discussion

In the present study we observe Neanderthal signatures in modern human mitochondrial genomes. Recombination test data here presented (bootscan analysis) shows that recombination does not explain the presence of all these 66 SNVs, or N-SNVs, in all modern mitochondrial haplogroups. The topic of recombination in mtDNA is hotly debated and evidence supporting it as well as evidence denying it are published [10,11]. It can be argued that recombination in human mtDNA is extremely rare, just sufficient for mtDNA to escape the Müller’s ratchet, but not enough to distinguish clear recombination blocks. We analyzed an alignment of 103 mitogenomes, which contained 918 polymorphic positions, by bootscan tests and PCA. The idea was to evaluate whether the presence of N-SNVs could be explained solely by homoplasy or recombination. We show that there are N-SNVs in present day human mtDNA that are completely absent in archaic modern Human mtDNA (41 mitogenomes). From our data we tested the possibility that rare recombination events might, at least in part, explain these results. Although an intense homoplasy rate could explain the N-SNVs in the D-loop, it is interesting to test if recombination can be associated with N-SNVs in coding regions, especially the nonsynonymous changes in positions 5460 (*ND2*, rs3021088), 7146 (*COX1*, rs372136420), 7650 (*COX2*, without rs, ACC>ATC), 9053 (*ATP6*, rs199646902), 13105 (*ND5*, rs2853501), 13276 (*ND5*, rs2853502) and 14178 (*ND6*, rs28357671). Our bootscan analyses suggest that all these N-SNVs could be explained by homoplasy in the last 40,000 years and that these changes occurred only in the present-day lineages and not in any of the archaic AMH mitogenomes analyzed.

The mitogenomes analyzed were from modern *Homo sapiens*, archaic AMH and Neanderthals as detailed above. Based on populational data Sykes (2001) estimated that a single observed change in comparative mitogenomics corresponds to 10,000 years of divergence. More modern estimates of the mitogenome clock, based on ancient DNA data, ranges from 2.14×10^−8^ to 2.74×10^−8^ substitutions per site per year [37–40], which gives approximately 4.14 substitutions in mitogenomes in 10,000 years. Therefore, according to populational estimates the 918 polymorphisms would have occurred in a period of 9.18 million years and the 66 N-SNVs for 660,000 years. If ancient DNA estimates are considered, the 918 polymorphisms would correspond to 2.21 million years of evolution and the 66 N-SNVs would have occurred in the last 159,420 years. Therefore all 66 N-SNVs are a product of random changes or simple homoplasy not observed in any of the 41 samples of archaic AMH. This suggests that homoplasy, parallel with Neanderthals, occurred only in the modern lineage, and none of those observed in archaic AMH. Tests for recombination using bootscan (Figure 6) indicate that in 11 positions the bootstrap support is significant for recombination which is not sufficient to explain all N-SNVs. It is important to notice that Posada and Crandall [41] explicitly tested how homoplasy could confound recombination tests and concluded that in extreme levels of rate variation (α=0.05) recombination tests would produce false positives which fits with mtDNA mutational load and among-site rate variation.

The analysis of Neanderthal mitochondrial genomes here presented revealed four derived amino acid changes that modern humans carry in the *COX2* gene as compared to Neanderthals and other Ape outgroups [42]. However, the same four amino acid changes can also be found in macaques which suggests that there is no need to invoke mitochondrial recombination, which is exceedingly rare, if present at all, or interbreeding events, to explain the presence of N-SNVs in modern human mitochondrial genomes and therefore recurrent mutations would be sufficient to explain N-SNVs. The divergence between the *H. sapiens* lineage and macaques lineage is 30.5 (26.9-36.4) million years [43] while the divergence we are dealing here is between 300,000 to 28,000 years. As is known, homoplasy significantly increase with long divergence times which produces the effect of long branch attraction in phylogenies [44]. For example, one of the mutations would be the macaques’ parallel mutation in *COX2* gene m.7650C>T. This mutation is found in Neanderthals (except Mezmaskaya), Denisovans, modern R0 haplogroup, Gorilla, Chimpanzee and Bonobo. It was not found in any archaic AMH samples. This pattern more likely suggests that it is highly conserved, inherited by early hominins from apes and secondarily lost in one Neanderthal lineage and in almost all *Homo sapiens* lineages, expect for the R0 haplogroup. The reappearance of this mutation in the very old R0 haplogroup has two possible explanations: (1) a back mutation reverting to exact the same ancestral condition instead of changing to any of the other three possible bases or (2) acquired by recombination *via* a Neanderthal female with introgression in one of the *Homo sapiens* lineages. In our study we present a bootstrap recombination analysis (bootscan) that shows a recombination point with bootstrap support in the region encompassing m.7650C>T (Figures 6A, 6E and 6F). Based on this test alone it seems that at least regarding this mutation a very rare recombination event could have happened although a back mutation cannot be completely excluded.

Another possible argument is that the Neanderthal signatures are in fact character states conserved since the last common ancestor of Neanderthals and present-day *Homo sapiens* (e.g. *Homo erectus* but this would not be consistent with the absence of these signatures in ancient *H. sapiens* mitochondrial genomes (Figure 1).

In regard to presence of N-SNVs in African haplogroups it is interesting to note that a back to Africa hypothesis has been proposed in which humans from Eurasia returned to Africa and impacted a wide range of sub-Saharan populations [45]. Our data suggest that Neanderthal signatures might be present in all major African haplogroups which is consistent with the “Back to Africa” contribution to the modern mitochondrial African pool. The preponderance of N-SNVs in the D-loop is observed mostly in African haplogroups. In Eurasian haplogroups we observe important changes in coding regions as demonstrated in Figures 2 and 3. Although our data suggest that convergent mutations explains the N-SNVs here observed, mitochondrial recombination is not so rare as to completely exclude it as a potential mechanism in human mtDNA mutational patterns [46,47].

Our observations suggest that crosses between AMH males and Neanderthal females left significantly less descendants than the reverse crosses (Neanderthal males and AMH females), which seems to be the dominant pattern. Although it has been generally accepted that recombination does not occur in the human mitochondrial genome, evidence of mitochondrial recombination has been reported [10,48]. A scenario with complete absence of recombination presents a problem to explain how the human mitochondrial genome would escape the Müller’s ratchet and therefore avoiding its predicted “genetic meltdown” [14]. It has been shown that even minimal recombination is sufficient to allow the escape from the Müller’s ratchet [49] and this could be the case regarding the human mitochondrial genome. For example, recombination has been simulated along a chromosome of 1000 *loci* to estimate the amounts of recombination required to halt Müller’s ratchet and the drift-catalyzed fixation of deleterious mutations. It has been found that for a population size of *N*<100, a recombination rate equivalent to one crossover per chromosome per 100 generations (10^−5^/locus/generation) countered Müller’s ratchet effectively. This is much lower than the minimum of one crossover per chromosome arm per generation that is assumed to occur in sexual taxa. A higher recombination rate of 10^−4^ can impede the selective interference that would otherwise enhance the fixation of deleterious mutations due to genetic drift [50].

Our data is compatible with a scenario in which the AMH-Neanderthal crosses occur in Europeans, East Asians and African lines of descent which is consistent with recent findings in nuclear genomes [51]. However, in the African haplogroups the crosses between AMH males and Neanderthal females would have a higher frequency than in European lines of descent, where the reverse crosses would be predominant. Based on the comparison of Neanderthal signatures in nuclear and mitochondrial genome haplogroups we hypothesize that the African lines of descent would have a higher female Neanderthal contribution whereas European lines of descent would have higher male Neanderthal contribution. The fact that AMH and Neanderthals crossed and produced fertile descendants is evidence that they belong to the same species [2]. Some authors propose that this suggests that *Homo sapiens* emerged independently in Africa, Europe and Asia [52]. The intercrosses of these three *Homo sapiens* subgroups, and other even deeper ancestors such as Denisovans, in their different proportions and specific signatures, would have produced the extant human genomes.

## 5. Conclusions

The analyses here presented suggest that Neanderthal genomic signatures might have been a product of convergent evolution due to homoplasy and that rare mtDNA recombination events cannot explain the presence of these signatures in modern mtDNA haplogroups. Although there is some evidence for mtDNA recombination, its weight in phylogenies and mutational patterns in comparative analysis remain controversial. Some authors contend that due to the high mutation rate in mtDNA, reverse compensatory mutations can be confounded with recombination. Our data is consistent with a scenario in which mtDNA recombination is not sufficient to support the contribution of Neanderthal mtDNA to modern human genomes, although rare recombination events might occur in human mtDNA to alleviate the effects of Müller rachet.

## Acknowledgments

The authors thank Prof. Dajiang Liu (Dept. Public Health Sciences, Penn State College of Medicine) and Prof. Rongling Wu (Center for Statistical Genetics, Penn State University) for critical reading of the manuscript. This work was supported by grants to M.R.S.B. from FAPESP, Brazil (2013/07838-0 and 2014/25602-6) and CNPq, Brazil (303905/2013-1). R.C.F received a CNPq postdoctoral fellowship (206445/2014-8) and C.R.R. a CAPES (Brazil) MSc fellowship.

## Author contributions

R.C.F. and C.R.R. planned and performed analyses. M.R.S.B. performed preliminary analysis and wrote the manuscript. R.C.F., C.R.R., J.R.B and M.R.S.B. discussed data and analyses and edited the manuscript.

## Data and materials availability

All data and files of analyses here presented are available upon request to the corresponding author.

